# Genomic analysis unveils important aspects of population structure, virulence, and antimicrobial resistance in *Klebsiella aerogenes*

**DOI:** 10.1101/581645

**Authors:** Hemanoel Passarelli-Araujo, Jussara K. Palmeiro, Kanhu C. Moharana, Francisnei Pedrosa-Silva, Libera M. Dalla-Costa, Thiago M. Venancio

## Abstract

*Klebsiella aerogenes* is an important pathogen in healthcare-associated infections. Nevertheless, in comparison to other clinically important pathogens, *K. aerogenes* population structure, genetic diversity, and pathogenicity remain poorly understood. Here, we elucidate *K. aerogenes* clonal complexes (CCs) and genomic features associated with resistance and virulence. We present a detailed description of the population structure of *K. aerogenes* based on 97 publicly available genomes by using both, multilocus sequence typing and single nucleotide polymorphisms extracted from core genome. We also assessed virulence and resistance profiles using VFDB and CARD, respectively. We show that *K. aerogenes* has an open pangenome and a large effective population size, which account for its high genomic diversity and support that negative selection prevents fixation of most deleterious alleles. The population is structured in at least ten CCs, including two novel ones identified here, CC9 and CC10. The repertoires of resistance genes comprise a high number of antibiotic efflux proteins as well as narrow and extended spectrum β-lactamases. Regarding the population structure, we identified two clusters based on virulence profile due to the presence of the toxin-encoding *clb* operon and the siderophore production genes, *irp* and *ybt.* Notably, CC3 comprises the majority of *K. aerogenes* isolates associated with hospital outbreaks, emphasizing the importance of its constant monitoring. Collectively, our results can be useful in the development of new therapeutic and surveillance strategies worldwide.

## INTRODUCTION

*Klebsiella aerogenes* is a Gram-negative, motile, rod-shaped bacterium, belonging to the *Enterobacteriales* order. It was formerly called as *Enterobacter aerogenes* according to Hormaeche and Edwards 1960 [1], but phylogenetic analysis showed that this bacterium is more closely related to *Klebsiella* than to *Enterobacter* species [2]. Widely considered an opportunistic pathogen, *K. aerogenes* is associated with nosocomial outbreaks, causing bloodstream, skin and soft tissue, respiratory, and urinary tract infections [3]. The emergence of *K. aerogenes* strains displaying multidrug resistance (MDR) has been associated with high mortality rates in patients from intensive care units [3].

Antibiotics have been extremely successful in treating several human infections and as a prevention strategy for different clinical interventions [4]. However, the widespread and often inappropriate use of antibiotics accelerates the emergence of resistant bacterial pathogens [5, 6]. Hospital outbreaks have already been related to *K. aerogenes* isolates carrying extended-spectrum beta-lactamase (ESBL) and carbapenemase genes [3, 7], resulting in the adoption of polymyxins as one of the last treatment options [8]. Antimicrobial resistance can also be associated to virulence factors [9], making the genomic analysis of multidrug resistant microorganisms a valuable tool to help preventing outbreaks and controlling the spread of multi-resistant and virulent strains.

A very effective means of investigating the evolution of a given population is through the characterization of its pangenome, which is the complete repertoire of genes found in several genomes (e.g. of different isolates) of a given species [10]. The pangenome is divided in the core genome, corresponding to genes common to all isolates; the accessory genome, comprising genes present in more than one, but not in all isolates and; unique (or strain-specific) genome, composed of genes found in a single genome. A pangenome can be defined as open or closed according to the capacity to acquire and use exogenous DNA [2, 11]. The open and closed pangenome concepts have also been linked with sympatric and allopatric lifestyles, respectively [2]. Each subdivision of the pangenome helps elucidate physiological aspects, lifestyle, genetic dynamics, pathogenicity, and population structure of a species [12, 13].

The ability to determine the genetic relatedness between bacterial pathogens is fundamental for epidemiological and evolutionary studies. This process can be facilitated when the desired population structure is defined by techniques such as Multilocus Sequence Typing (MLST), which is available for several species [14]. This technique is based on the analysis of single-nucleotide polymorphisms (SNPs) within housekeeping genes, with each locus receiving a different allele number. Each unique allelic profile (or genotype) is a sequence type (ST), which often defines a clone or strain and can be linked to other STs to form Clonal Complexes (CCs) [15]. The linkage of STs to form a CC is based on the number of differences between allelic profiles. For example, genotypes which differ at only one locus are single-locus variants (SLVs), those with two different *loci* are double-locus variants (DLVs). It is also possible to determine lineage linkages with core genome MLST (cgMLST), which has greater resolution than classical MLST [16]. The ST and CC information can be used to uncover relationships between closely-related isolates and to elucidate the population distribution across different geographic regions [17], allowing the elaboration of surveillance strategies to prevent the dissemination of specific strains.

The balance between selection and drift plays important roles in shaping bacterial genomes and are often studied by estimating the selection coefficient and the effective population size (*N*_*e*_) [18, 19]. Due to several technical and biological reasons (e.g. the existence of truly neutral sites in bacterial genomes), it has proven hard to estimate *N*_*e*_ based on neutral expectations [19]. An interesting alternative that has been successfully applied in bacterial genomics is the use of substitution rates at non-synonymous and synonymous sites (*d*_N_/*d*_S_) to estimate *N*_e_. Because *d*_N_/*d*_S_ ratios directly reflect the selection strength [20] and negatively correlates with *N*_*e*_, it can be useful to understand the evolutionary dynamics of different bacterial populations, including those of clinically important pathogens [21].

*K. aerogenes* was included in the ESKAPE group [22, 23] of important pathogens typically associated with antimicrobial resistance. However, in contrast to other ESKAPE species (e.g. *Staphylococcus aureus* and *Pseudomonas aeruginosa*), the pangenome and population structure of *K. aerogenes* have not been extensively explored. Here, we report a comprehensive comparative genomic analysis of *K. aerogenes*, which allowed us to uncover important features of its pangenome, selective pressure acting in the species, population structure, resistance, and virulence profiles.

## RESULTS AND DISCUSSION

### GENOME ASSEMBLY AND PREVALENCE OF PLASMIDS AND BACTERIOPHAGES

We systematically obtained raw sequencing reads, assembled genomes, and predicted genes of 91 *K. aerogenes* strains with publicly available data. This dataset was were supplemented with six reference genomes (that were not assembled and annotated *de novo*) (see methods for details). All the genomes showed at least 95% of genome completeness according to BUSCO [24]. *K. aerogenes* has an average genome size of 5,281,261 ± 194,500 bp and 4984 ± 219 genes. We found an average %GC of 54.95 ± 0.17 in *K. aerogenes*, which is within the expected range for *Klebsiella* [21] (Supplementary Table 1). ANI analysis revealed that all the isolates have over 95% genomic identity, confirming that all the strains belong to the same species.

**Table 1.** Prevalence of plasmids in *K. aerogenes* genomes. The proportion of isolates with plasmids containing resistance and virulence genes and the median number (min – max) are indicated. “Untypeable” plasmids are those for which a replicon was not identified using PlasmidFinder.

| Replicon | Number of isolates | Antimicrobial resistance genes |  | Virulence genes |  |
| --- | --- | --- | --- | --- | --- |
|  |  | Proportion (%) | Median number | Proportion (%) | Median number |
| Col | 4 | 0 (0%) | 0-0 | 0 (0%) | 0-0 |
| Col156 | 4 | 0 (0%) | 0-0 | 1 (25%) | 1-1 |
| ColpVC | 2 | 0 (0%) | 0-0 | 0 (0%) | 0-0 |
| ColRNAI | 49 | 1 (2%) | 2-2 | 0 (0%) | 0-0 |
| IncA/C2 | 5 | 4 (80%) | 7-12 | 0 (0%) | 0-0 |
| IncB/O/K/Z | 1 | 1 (100%) | 1-1 | 1 (100%) | 1-1 |
| IncFIA | 6 | 5 (83%) | 1-1 | 6 (100%) | 1-1 |
| IncFIB | 21 | 7 (33%) | 1-7 | 11 (52%) | 1-7 |
| IncFII | 28 | 9 (32%) | 1-7 | 14 (50%) | 1-4 |
| IncHI2 | 2 | 1 (50%) | 3-3 | 1 (50%) | 1-1 |
| IncL/M | 3 | 0 (0%) | 0-0 | 0 (0%) | 0-0 |
| IncN | 3 | 2 (66%) | 1-8 | 3 (100%) | 1-1 |
| IncQ1 | 2 | 2 (100%) | 3-3 | 0 (0%) | 0-0 |
| IncR | 4 | 4 (100%) | 1-1 | 4 (100%) | 1-1 |
| IncX3 | 5 | 2 (40%) | 2-2 | 3 (60%) | 1-1 |
| IncX4 | 1 | 0 (0%) | 0-0 | 0 (0%) | 0-0 |
| pENTAS02 | 1 | 1 (100%) | 14-14 | 1 (100%) | 19-19 |
| Rep | 1 | 1 (100%) | 1-1 | 0 (0%) | 0-0 |
| Untypeable | 28 | 12 (42%) | 1-9 | 2 (7%) | 4-4 |

We inspected the assembled genomes for possible plasmids based on plasmid replicons, identity, and coverage with publicly available plasmid sequences (Supplementary table 3). Plasmid replicons or fragments with high identity and coverage were found in 70.1% of the isolates, ranging from one to 10 plasmids per strain with most isolates harboring two plasmids. The most abundant replicon was ColRNAI (Table 1), which is related to plasmids carrying genes related with toxin-antitoxin system [25]. The second most prevalent plasmids were those with IncF replicons, which are low copy-number plasmids associated with the dissemination of resistance genes, such as *bla*_CTX-M_ and *acc(6’)-Ib-cr* [26].

Although abundant in the population, the ColRNAI replicon type showed a low proportion of resistance and virulence genes. Plasmids with the IncA/C2 incompatibility group were found in EA1509, G7, D2, and C10, which are strains associated with hospital outbreaks. These plasmids harbored a high number (i.e. seven to 12) of resistance genes (Table 1), but lacked virulence genes. The strain with the highest number of virulence determinants in plasmids (19 genes) was SMART_1248, which was collected from an old-male urine sample from a surveillance study [27]. Isolates with uncharacterized plasmid replicons were also recovered (Table 1).

By using the PHASTER database (Supplementary table 4), we found a variety of 72 different bacteriophages, with frequencies ranging from one to 11 in *K. aerogenes* genomes. Importantly, we recovered ST-specific phages, such as the Enterobacteria phage 933W (NC_000924), found in 91.66% of ST4 isolates. The 933W phage belongs to a group that encodes the Shiga-like toxin and is associated with *E. coli* infections [28]. Since most reports of this phage are in *E. coli* [29], its presence in *K. aerogenes* ST4 may indicate a horizontal gene transfer (HGT) event that carried a toxin gene to *K. aerogenes*.

### PANGENOME ANALYSIS

We found that *K. aerogenes* has an open pangenome with a total of 18,268 gene families (Figure 1a), indicating a high level of genetic diversity and allowing us to predict that many more additional gene families will be detected as new genomes are sequenced. The open pangenome of *K. aerogenes* is also in line with its sympatric lifestyle [2]. That is, due to living in communities with a high diversity of microorganisms, it tends to acquire novel genes by HGT. A previous study estimated the *K. aerogenes N*_*e*_ in 338,041,563.813 [21]. As discussed above, such high *N*_*e*_ is an evidence of strong purifying selection, favoring the retention of numerous accessory genes (including those acquired from HGT) that could ultimately account for the open pangenome of this species.

**Figure 1.**
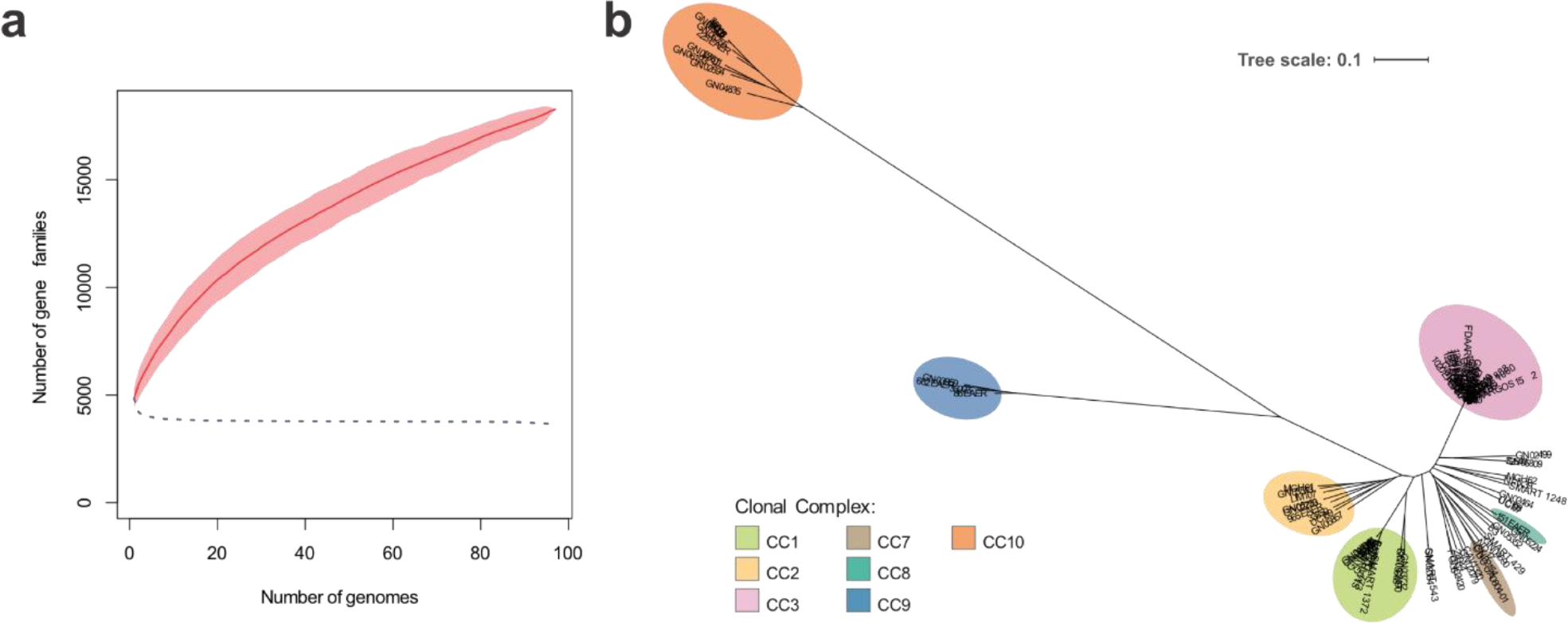
Pangenome and phylogenetic analysis of *Klebsiella aerogenes*. (a) Growth of the core genome (dotted line), pangenome (red solid line) and its confidence intervals (light red area). This plot was generated using the vegan package 2.5-2 [70]. The cumulative curve is characteristic of an open pangenome; (b) Maximum likelihood phylogenetic reconstruction performed with 264,006 SNPs identified in the core genome. Colors indicate clonal complexes (CCs) determined by cgMLST. Isolates outside of colored circles were not assigned to any CC.

The *K. aerogenes* core genome has a total of 3,766 genes (Figure 1a). In other words, 20.61% of the pangenome is associated with intrinsic physiological traits that evolve vertically and under strong purifying selection due to their essentiality [30, 31]. We used 264,006 SNPs in the core genome to perform a maximum likelihood phylogenetic reconstruction (Figure 1b), which showed that some clusters are distant from the main part of the population, supporting the intra-specific diversity of *K. aerogenes*.

In order to estimate the efficiency of selection in *K. aerogenes*, we calculated π_syn_= 0.05 and *d*_*N*_/*d*_*S*_ = 0.04, which corroborate previous findings and explain the high *N*_*e*_ discussed above. In fact, *d*_*N*_/*d*_*S*_ has been shown to be a robust proxy of *N*_*e*_[19]. Because the *d*_*N*_/*d*_*S*_ is quantitatively different when individuals from the same species are analyzed, the classical signatures of *d*_*N*_/*d*_*S*_ < 1 as indicating weak negative selection in divergent populations have a different interpretation in intraspecific studies [20]. According to the ranges determined in a previous study [20], our results indicate that *K. aerogenes* evolves under a strong negative selection.

The accessory genome is composed by medium- and low-frequency genes. A total of 1,549 genes were found between 15% and 94% of the population and 12,953 genes were present in ≤15% of the genomes. The abundance of rare genes further reflects the genome plasticity of *K. aerogenes* and probably play roles in niche adaptation and ecological dynamics, in particular because of their localization in genomic islands or plasmids containing transposases and integrases [32, 33]. This finding is also in line with our results presented above on the prevalence of plasmids harboring toxin-antitoxin systems (>50%), which that likely allow *K. aerogenes* to exploit new niches by increasing its competitiveness, fueling the emergence of new subpopulations.

### POPULATION STRUCTURE ANALYSIS UNCOVERS NEW *K. aerogenes* CLONAL COMPLEXES

A *K. aerogenes* MLST scheme was published in December 2017. From the 140 reported STs, 25.7% belong to a known CC. We locally updated this database with 33 new STs and two new CCs, which we named CC9 and CC10 (Supplementary Table 5). According to our updated MLST analysis of sequenced isolates, at least 10 CCs compose the *K. aerogenes* population structure (Figure 2): CC1 (n = 16); CC2 (n = 8); CC3 (n = 33); CC5 (n = 5); CC7 (n = 2); CC8 (n = 1); CC9 (n = 4); CC10 (n = 11). Further, it was not possible to assign a CC to 19 isolates, which were called NOCC. We were unable to recover isolates from CC4 and CC6. Importantly, ANI analysis confirmed the genetic distance from CC10 and CC9 to other CCs (Supplementary figure 1).

**Figure 2.**
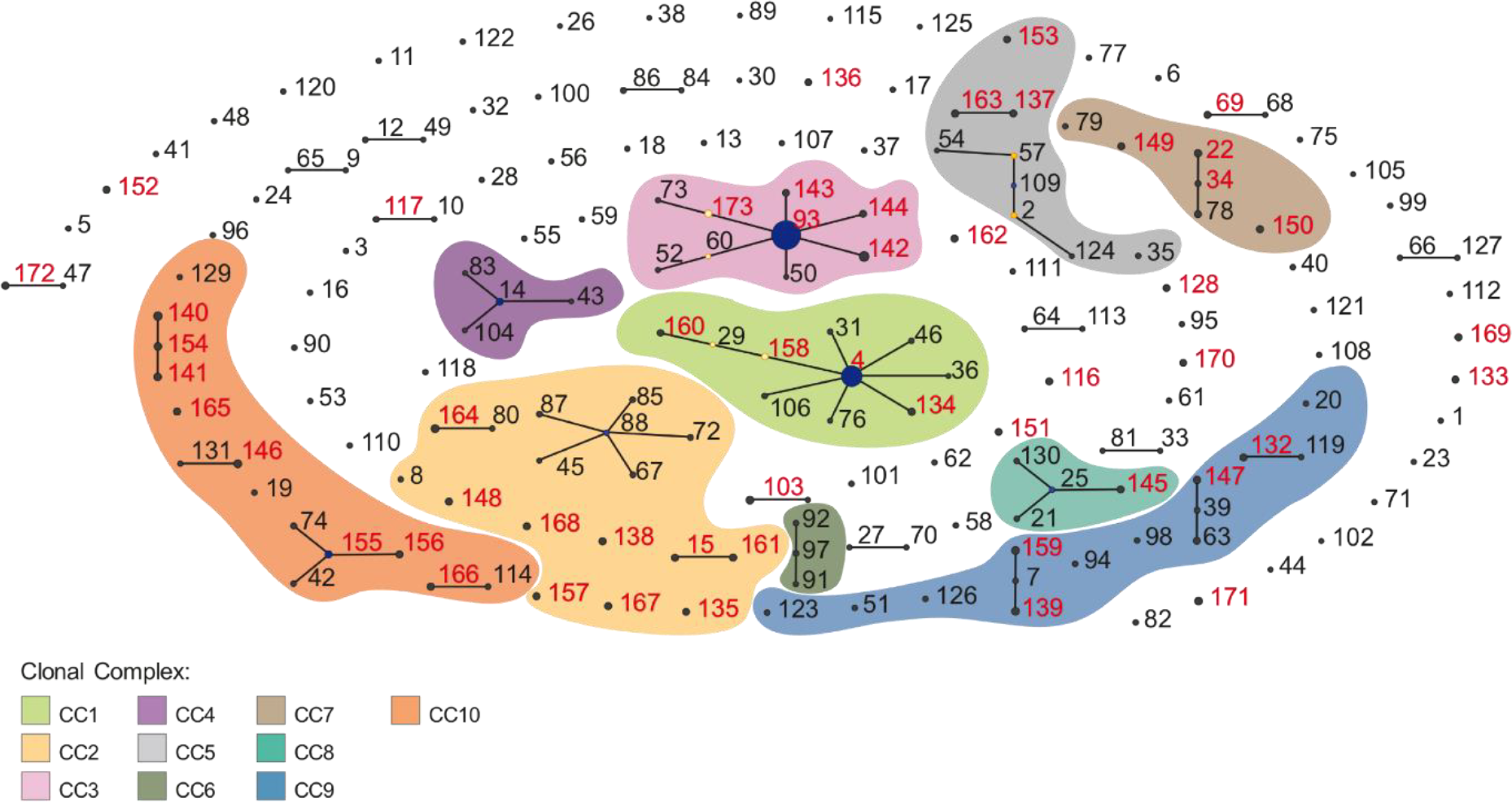
Population structure of *Klebsiella aerogenes* based on MLST. Black labels represent STs available in the database absent in our study, while red labels refer to STs analyzed here, including novel ones. SLVs and DLVs are linked by edges, resulting the formation of 10 clonal complexes (CCs). Yellow dots represent secondary SLVs, which are SLVs of SLVs. Results from the cgMLST analysis were used to improve the resolution of CC determination, accounting for the assignment of STs to CCs without edges. Blue dots represent founder STs (ST4, ST14, ST25, ST88, ST93, ST109, ST155) and help understand the origin and diversification of each CC. Founder STs were not predicted for CC6 and CC7. Dot sizes are proportional to ST frequencies. We were unable to recover samples from CC4 and CC6. The two new CCs identified here (CC9 and CC10) are also represented.

To increase the resolution arising from the use of few genes in the traditional MLST technique, we also used the SNPs from the core genome to perform a cgMLST structure (Figure 1b). This analysis improved the discriminative power of each CC and allowed us to better define the population structure of *K. aerogenes*. Importantly, the same population structure was inferred with both methods, supporting the presence of at least 10 CCs.

The most prevalent STs were ST93 and ST4, with ~27% and ~13% of the isolates, respectively. A large number of STs were singletons, as they had no SLVs or DLVs to form groups (Figure 2). Importantly, the strains used in our study (Figure 2, red dots) contributed to a more precise characterization of some CCs (e.g. CC3), emphasizing the need for a constant MLST characterization and ST/CC assignment of new isolates. Finally, upon submission of our manuscript, we have shared the information on the new CCs identified here with the pubMLST database team.

In addition to its importance in pathogen surveillance, CC determination is also critical in evolutionary studies. For example, CC3 is the one with the greatest number of isolates and has ST93 as its founder genotype (Figure 2). The term founder ST derives from population genetics to describe situations where low population diversity gives rise to a genotypically related group. Thus, the founder ST helps understand the evolutionary trajectory of a given CC (Figure 2). ST155 is the predicted founder of CC10, which has GN04835 as its most basal member (Figure 1b), being associated with its diversification. This pattern is also observed for the other CCs, allowing a better understanding of the diverse resistance and virulence profiles found in the *K. aerogenes* population, as we show in the next sections

### RESISTANCE GENES AND THEIR DISTRIBUION ACROSS *K. aerogenes* CLONAL COMPLEXES

We found a total of 94 and 95 antimicrobial resistance genes the *K. aerogenes* core and accessory genomes, respectively; these gene sets were called core and acquired resistome, respectively (Supplementary table 6). Regarding the core resistome, we confirmed CMY-108 as the chromosomal constitutive AmpC β-lactamase, whose overexpression leads to cephalosporin resistance [34]. Another core gene conferring resistance against β-lactams is that encoding the outer membrane porin OmpK37, originally associated with lower membrane permeability to the antibiotic in *K. pneumoniae* [35]. Other core genes were predicted to confer resistance against aminocoumarin, aminoglycosides, macrolides, fosfomycin, fluoroquinolones, sulfonamides and tetracyclines (Figure 3c, Supplementary table 6).

**Figure 3.**
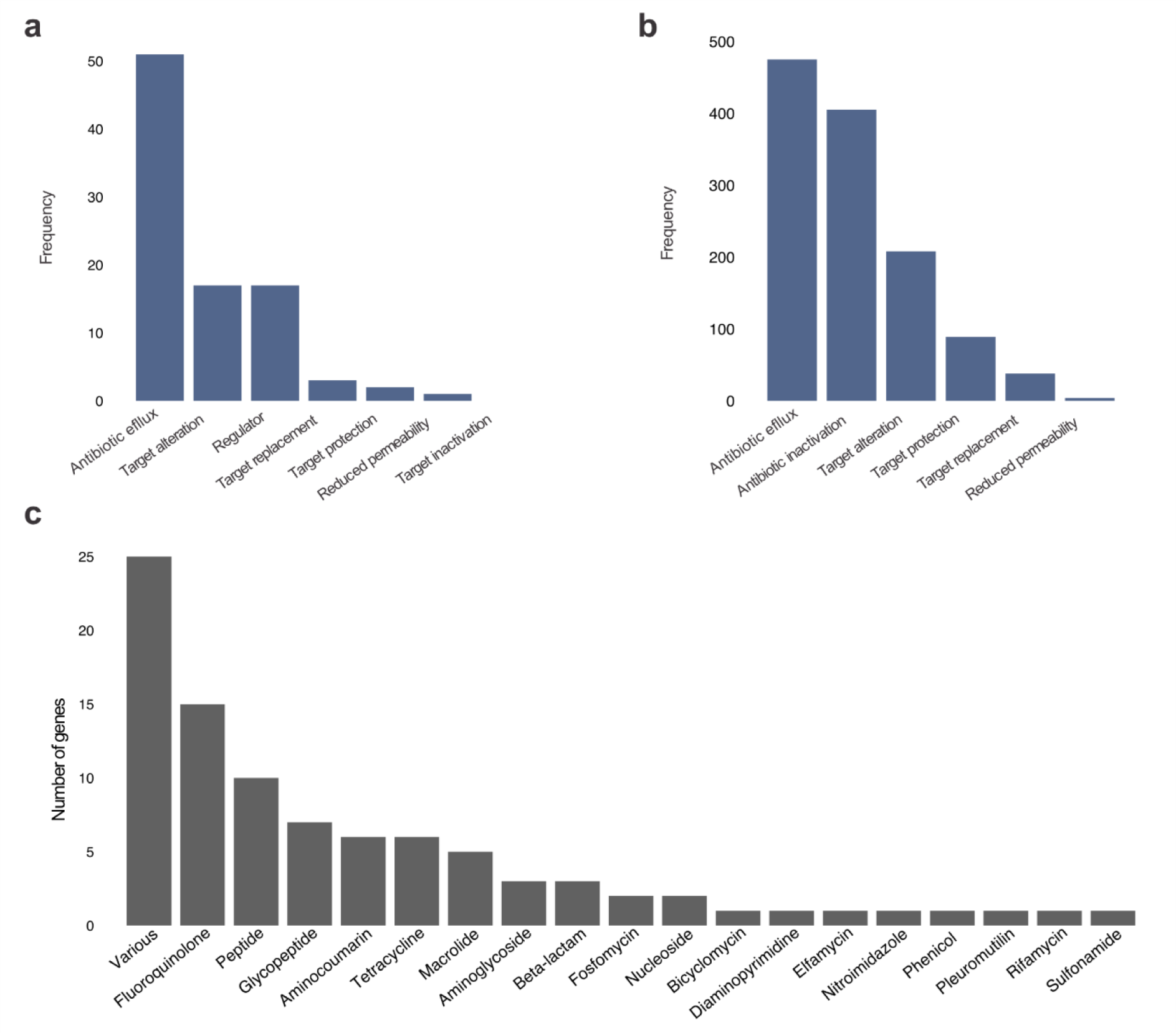
Main antibiotic resistance mechanisms. (a) Core resistome and (b) acquired resistome. In both cases, the main mechanism is antibiotic efflux. However, the core resistome has multiple genes encoding regulators and antibiotic target modifiers, whereas the acquired resistome harbors a high number of antibiotic inactivating proteins. (c) Antibiotic classes to which core genes confer resistance to. Genes associated with resistance against different antibiotic classes were classified as “various”. Many of these genes encode efflux pumps and regulators.

A range of MDR efflux pumps (e.g AcrAB-TolC, AcrD, EmrAB-TolC, MacAB-TolC, MdtABC, OqxAB, RosAB) compose the main resistance mechanism encoded by the core resistome (Figure 3a), potentially conferring resistance against different antibiotic classes, such as fluoroquinolones and macrolides. Indeed, Gram-negative bacterial pathogens exhibit relatively high resistance to macrolides due to the high number of chromosomal-encoded MDR efflux pumps [4]. Further, AcrAB-TolC inactivation by inhibitors induced the susceptibility of *K. aerogenes* to macrolides [36]. Similarly, the inhibition of some MDR efflux pumps rendered *Pseudomonas aeruginosa* susceptible to fluoroquinolones [37]. We hypothesize that the characterization of the intrinsic resistome can aid in the prospection of new antibiotic targets and in the identification of potential drug combinations.

In addition to proteins directly involved with resistance, we also identified regulators that control the expression of resistance genes. This gene set includes: BaeRS, the two-component regulatory system associated with the transcriptional activation of MdtABC efflux pumps [38]; the stress-sensor MarA that promotes the AcrAB-TolC expression and RarA, a positive regulator of AcrAB and MarA in *K. aerogenes* [39, 40] and; EmrR, a negative regulator of the EmrAB-TolC efflux pump [41]. These results demonstrate that the inactivation of specific regulatory systems in combination with current antibiotics could produce pleiotropic effects that might increase antibiotic efficacy and even allow the resurrection of obsolete antibiotics.

As opposed to the core resistome, acquired resistance genes are often found associated with mobile elements, suggesting their origin via HGT. The main resistance mechanism found in the *K. aerogenes* acquired resistome involves a large number of antibiotic efflux pumps and enzymes associated with antibiotic inactivation (Figure 3b). The acquired resistome also comprises major resistance determinants against aminoglycosides (*aac(3)*-IIa, *aac*(6’)-*Ib*, *aadA*, *aph*(6’)-*Ic*, *rmt*G), β-lactams (*bla*_TEM-1, 24, 215_, *bla*_SHV-66,_ *bla*_OXA-1, 2, 9, 48_, *bla*_CTX-M-2, 15, 59_, *bla*_CMY-108,_ *bla*_KPC-1, 3_, *bla*_NDM-6_ and *bla*_OXY-2-10_), diaminopyrimidine (*dfrA*), fluoroquinolones (*qnrB*), fosfomycin (*fosA*), macrolides (*rlmA, mphA, chrB*), sulfonamide (*sul1, sul2*) and tetracycline (*tet(A/D/G)*) (Supplementary table 6). By analyzing the frequency of each gene based on antibiotic classes that they confer resistance to (Figure 4), we found, for example, that CC10 tend to be more resistant against aminoglycosides.

**Figure 4.**
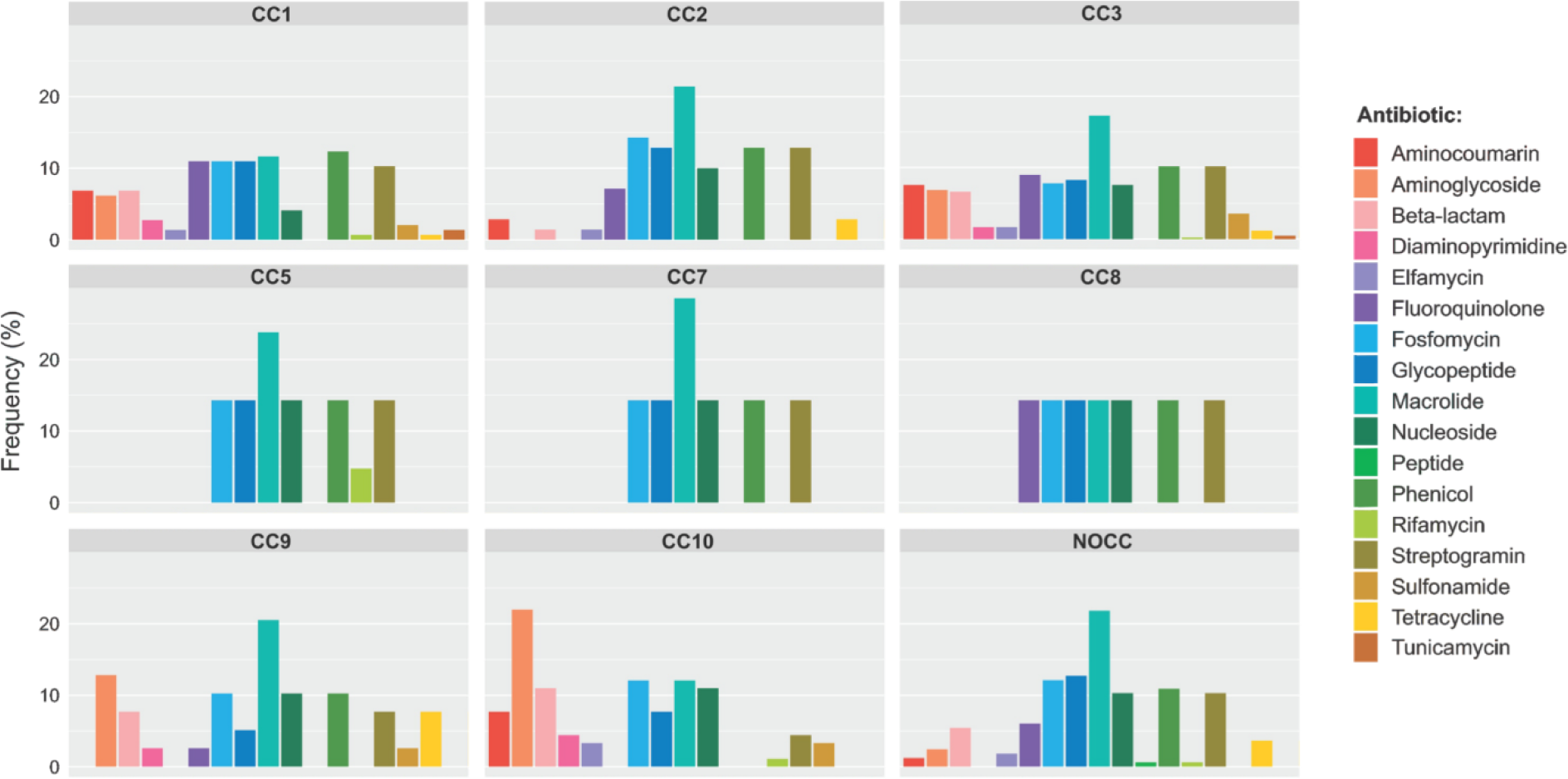
Relative acquired resistome size per clonal complex. In each box, we show the percentage of genes associated with the resistance of each class of antibiotics. It is clear from these results that aminoglycoside resistance genes are prevalent in CC10 (characterized in this study) and that macrolide resistance is widespread in almost all CCs. It is important to mention that CC5, CC7 and CC8 have fewer isolates. For better visualization, the “various” category from figure 3 is not represented here.

### VIRULENCE PROFILES ALLOW THE SEPARATION OF *K. aerogenes* STRAINS IN TWO CLUSTERS

We identified several genes encoding virulence factors in *K. aerogenes*. Our reference virulence database (see methods for details) comprises genes involved in biofilm formation, adherence, immune evasion, iron acquisition, allantoin utilization, regulators, secretion systems, serum resistance and toxin production. More than 70% of the reference genes were detected in at least one *K. aerogenes* isolate (Supplementary table 7). Virulence factors shared by all isolates included the *fimA-K* operon, involved in adherence to human mucosal or epithelial surfaces [42]; *ent*, *fep* and *iro*, which encode enterobactin and salmochelin siderophores [43, 44]; *rcsAB*, regulators of mucoid phenotype A [45], among others (Supplementary table 7). We also found intrinsic genes encoding toxins related to *K. aerogenes* physiology during infection, such as colibactin (ClbD), enterotoxin (SenB) and microcin (MceG and MceH), which could be explored as novel drug targets. On the other hand, virulence factors that were not common for all isolates were highly variable in terms of frequency across genomes (Figure 6; Supplementary table 7,).

The evolution of antibiotic resistance and virulence factors are often associated [9, 46]. Thus, we compared the prevalence and distribution of these genes across the whole population (Figure 5a). Most of the isolates have a relatively low frequency of resistance genes, while virulence factors display a bimodal distribution. When CCs are compared individually, there is a variation between the number of resistance and virulence genes (Figure 5b). For example, CC3 has a higher number of virulence genes when compared to resistance genes. It is important to note that more than 50% of the CC3 isolates derive from hospital outbreaks [2, 7, 47–49]. We hypothesize that, due to the sympatric lifestyle of *K. aerogenes* (i.e. open pangenome) and its occurrence in highly competitive environments, more virulent strains tend to increase in frequency in the population as a strategy to deplete the host resources faster increasing the transmission rate, as stated by the trade-off hypothesis [50, 51].

**Figure 5.**
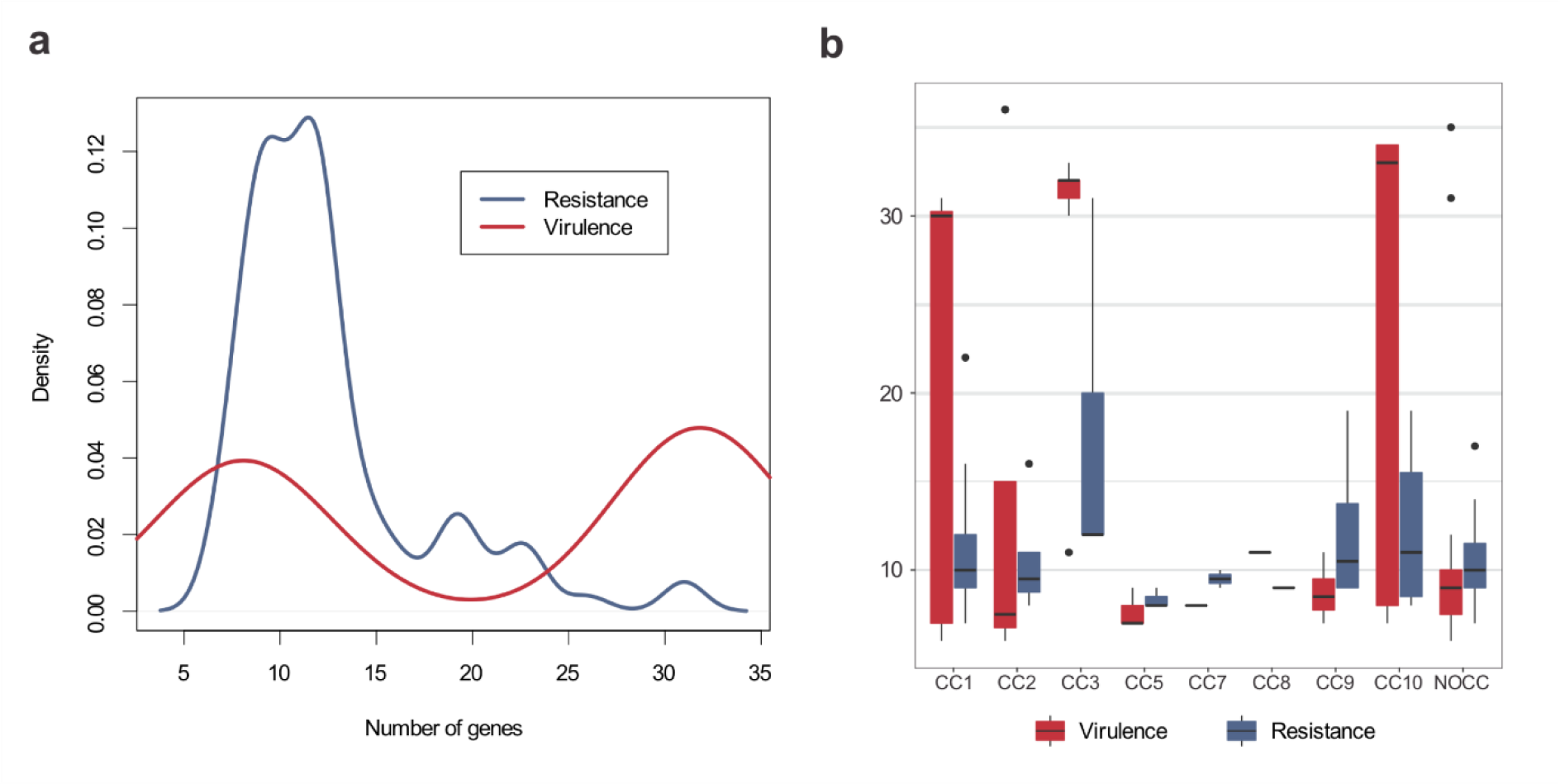
Comparison between acquired resistance and virulence profiles. (a) Density plot of the number of resistance (blue) and virulence (red) genes per strain. (b) Distribution of acquired resistance and virulence genes across clonal complexes (CCs). It is important to mention that CC5, CC7 and CC8 have fewer isolates.

**Figure 6.**
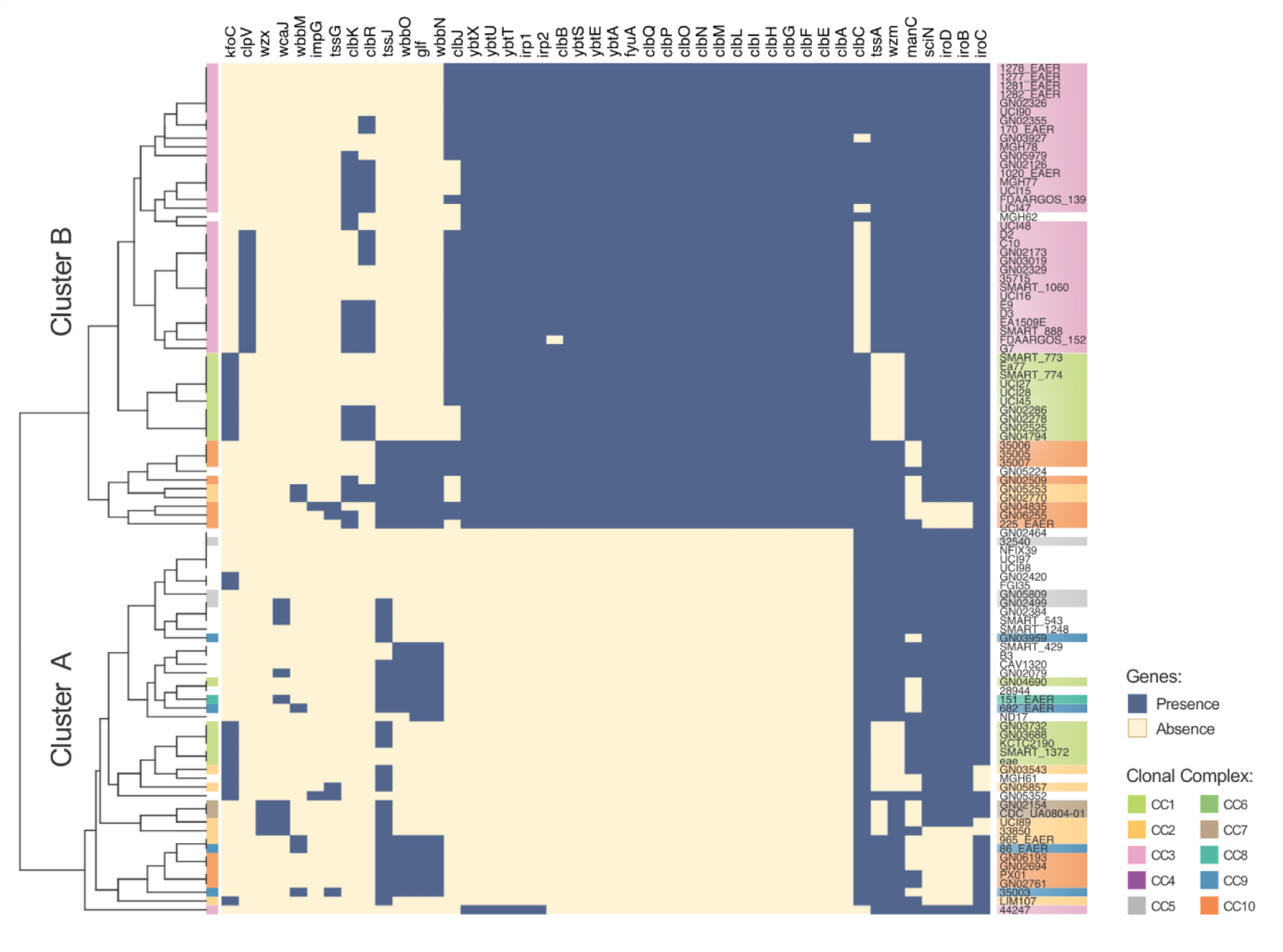
Prevalence of acquired virulence factors. Columns and rows represent virulence factors and strains, respectively. Strains are colored according to their CC. Two clusters can be easily identified: cluster A (less virulent) and cluster B (more virulent). Cluster B is typified by the colibactin *clbA-R* operon, and by *ybt* and *irp* operons, which are associated with siderophore production in *Yersinia* [71]. All CC3 strains (except 44247) belong to cluster B and are associated with hospital outbreaks.

To investigate whether the bimodal distribution of virulence genes (Figure 5a) was restricted to a specific CC, we built a presence/absence matrix to evaluate virulence profiles. The results indicate a clear separation between more and less virulent strains (Figure 6), referred here as Cluster A and B, respectively. Cluster B is basically defined by the presence of the *clb* operon, responsible for the production of colibactin, a genotoxic hybrid polyketide-nonribosomal peptide encoded by *psk* islands [52] and; *ybt* and *irp*, related to the production of yersiniabactins siderophores. Ybt is the most common virulence factor associated with human *K. pneumoniae* infections [53]. A very recent study also reported the high prevalence of yersiniabactins and colibactin in *K. aerogenes* ST4 and ST93 [54]. In addition, the most virulent cluster comprises isolates from CC1, CC2, CC3 (all isolates except 44247), and the previously uncharacterized CC10. These findings further highlight the importance of rapid and robust CC identification in the definition of different measures to slow the spread of virulent clones.

Altogether, our results support the capacity to distinguish the virulence profiles between CCs. We determined the population structure of *K. aerogenes*, including the identification of new STs and CCs. This update in the MLST scheme is useful for epidemiological and evolutionary studies. The next steps to advance the understanding of *K. aerogenes* could involve a more in-depth study of a greater number of isolates of each CC, as well as the characterization of new STs and CCs. In this context, sequencing samples from a wider range of locations is of great importance. Given the open pangenome and diversity of *K. aerogenes* isolates, it is likely that several novel STs and CCs will be identified as samples from different countries are sequenced. Our work provides new insights into key biological aspects (e.g. virulence and resistance) of *K. aerogenes*, as well as some interesting perspectives for the investigation of new drug targets, diagnostic and surveillance strategies.

## MATERIALS AND METHODS

### ISOLATES, GENOME ASSEMBLY, AND GENE PREDICTION

We conducted this study with 91 isolates of *K. aerogenes* with whole-genome sequencing reads available on Genbank on December 2017 (Supplementary Table 1). Sequencing quality was assessed with FastQC version 0.11.5 (https://www.bioinformatics.babraham.ac.uk/projects/fastqc/) and low-quality reads and adaptors removed with Trimmomatic 0.35 [55]. Non-reference genomes were assembled using SPAdes 3.11.1 [56] and scaffolds obtained with SSPACE 3.0 [57]; scaffolds with less than 500 bp were removed. These *de novo* assembled genomes were supplemented with six reference genomes that were directly used. Genome completeness was assessed with BUSCO 3.0 [24], using *Enterobacteriales* as a reference dataset. ANI (Average Nucleotide Identity) was computed with pyani 0.2.0 [58].

Plasmids were detected with the plasmidSPAdes pipeline [59]. Putative plasmid scaffolds were evaluated using plasmidFinder [60], which contains replicons of several complete plasmids belonging to major incompatibility (Inc) groups of Enterobacteriaceae species. In order to identify plasmids lacking replicon information, we also searched all plasmids of the GenBank database (January 2017) using BLASTN [61] with 70%, 50% and 50% of identity, query coverage, and subject coverage thresholds, respectively. ORFs from chromosomal and plasmid sequences were predicted and annotated with Prokka 1.12 [62]. Bacteriophage signatures were assessed with PHASTER [63].

### PANGENOME ANALYSIS AND PHYLOGENY

The *K. aerogenes* pangenome was inferred using Roary 3.6 [64]. Core genes were defined as those present in more than 95% of the isolates. Core gene sequences were aligned with MAFFT v7.271 [65]. SNPs were extracted from the core genome alignment and used for maximum likelihood phylogenetic reconstructions with RAxML [66], using the general time reversible (GTR) model and gamma correction specifying the use of only variable sites as input. One thousand bootstraps replicates were generated to assess the significance of internal nodes. The estimation of π_syn_ and *d*_*N*_/*d*_*S*_ were conducted with Mega X [67].

### MLST ANALYSIS

To determine the ST of each isolate, the pubMLST database [14] of *K. aerogenes* was used to identify SNPs in seven housekeeping *loci*: *dnaA, fusA, gyrB, leuS, pryG, rplB*, and *rpoB.* CC identification was performed with eBURST [17] to link SLVs and DLVs using 1,000 bootstrap resamplings. In addition, to allow a more precise distinction between CCs, we inferred the cgMLST scheme using the core-genome SNP phylogenetic tree.

### VIRULENCE AND ANTIMICROBIAL RESISTANCE GENES

The repertoire of resistance genes (i.e. the resistome) was inferred using the Comprehensive Antimicrobial Resistance Database (CARD) database (version 1.1.8) [68]. Virulence factors were analyzed using Virulence Factor Database (VFDB; accessed on October 2018) [69] and other virulence factors directly obtained from the literature (Supplementary Table 2). All predicted protein sequences were compared with these two databases with BLASTP [61] using minimum similarity and query coverage thresholds of 50% and 60%, respectively. Proteins that reached this threshold but were at least two-fold longer than the reference sequences were removed. In all methods mentioned in the materials and methods section, default parameters were used unless stated otherwise.

## ACKNOWLEDGEMENTS

This work was supported by the Fundação Carlos Chagas Filho de Amparo à Pesquisa do Estado do Rio de Janeiro and by Conselho Nacional de Desenvolvimento Científico e Tecnológico (CNP_q_). HP-A fellowship was funded by UENF. KCM and FP-S postgraduate fellowships were funded by Coordenação de Aperfeiçoamento de Pessoal de Nível Superior (CAPES; Finance Code 001).

---

**Supplementary Figure 1. Average nucleotide identity analysis of the 97 isolates used in this study.** Pairwise comparison indicating major groups in *K. aerogenes* based on nucleotide identity.

