## Supplementary figures and images for "Genomic analysis unveils important aspects of population structure, virulence, and antimicrobial resistance in *Klebsiella aerogenes*"

### Sup. Fig. 1

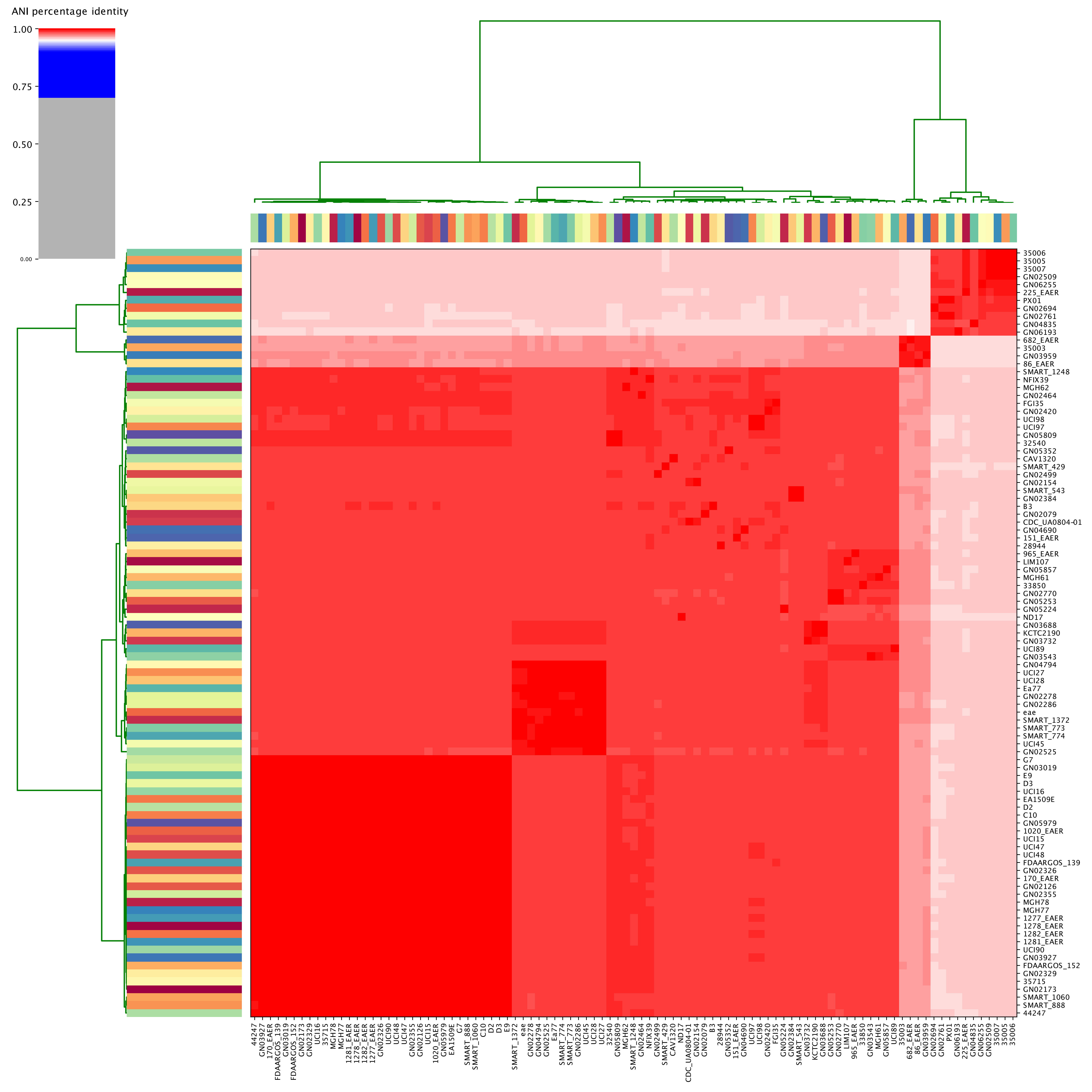
